# Emulating the influence of exoskeleton stiffness on primary afferent feedback in rat isolated muscle-tendon unit

**DOI:** 10.1101/2025.11.03.686366

**Authors:** Amro A. Alshareef, Paul Nardelli, Surabhi N. Simha, Timothy C. Cope, Lena H. Ting, Gregory S. Sawicki

**Affiliations:** Woodruff School of Mechanical Engineering, Georgia Institute of Technology, Atlanta, Georgia, USA; School of Biological Sciences, Georgia Institute of Technology, Atlanta, Georgia, USA; Coulter Department of Biomedical Engineering, Emory University and Georgia Institute of Technology, Atlanta, Georgia, USA; Department of Rehabilitation Medicine, Emory University, Atlanta, Georgia, USA

**Keywords:** sensory feedback, exoskeletons, proprioception

## Abstract

Exoskeletons assist and augment movement, but their effects on proprioceptive feedback remain poorly understood due to challenges in making direct measures of sensory signals in humans. Here, we leveraged a benchtop animal model to begin to explore how mechanical context akin to an elastic exoskeleton operating on a human lower limb joint may influence primary muscle spindle firing. In an anesthetized rat preparation, we applied controlled stretches to the medial gastrocnemius with engineered springs (0–0.5 N/mm) attached in parallel to the muscle-tendon unit (MTU) while modulating muscle activation to maintain overall system stiffness. Fascicle length was measured with sonomicrometry, force and MTU length with a servo motor, and spindle instantaneous firing rate (IFR) using dorsal root recordings. Trading off increases in parallel ‘exoskeleton’ stiffness with reductions in muscle activation decreased biological muscle force (3.1 ± 0.6 N to 1.6 ± 0.6 N, p < 0.001) and stiffness (4.4 ± 1.5 N/mm to 2.3 ± 1.3 N/mm, p < 0.01), and increased fascicle length (7.9 ± 1.3 mm to 8.3 ± 1.5 mm, p < 0.005). We found significant correlations between spindle firing and each independent muscle fascicle kinematic and kinetic factor we investigated (p < 0.005). Thus, parallel stiffness emulating a passive elastic exoskeleton modifies muscle fascicle dynamics but does not alter spindle firing, possibly due to internal trade-offs in the salient muscle fascicle kinetic and kinematic features that drive spindle behavior. Leveraging in situ experimental frameworks that enable monitoring of primary afferent feedback during active contractions in complex mechanical contexts such as added parallel stiffness can provide a window into the effects of wearable devices on underlying sensory systems.

**Summary Statement:** In an *in-situ* rat model replicating human exoskeleton dynamics, we show that increased exoskeleton stiffness alters muscle fascicle mechanics without changing muscle spindle firing rates.

## Introduction

Exoskeletons and exo-suits are emerging as a viable technology for restoring, assisting or augmenting human movement. Despite this, little focus has been placed on how these exoskeleton assistance profiles influence the user’s internal sensorimotor control loop. In recent years, exoskeletons have been developed to enhance performance during unsteady locomotion tasks that require dynamic force response, such as maneuvering to change direction or stabilizing to recover balance (Bayón et al., 2022; Beck et al., 2023; Bywater et al., 2023; King et al., 2022; Monaco et al., 2010; Robertson et al., 2017). Indeed, powered ankle exoskeletons can enhance standing balance recovery by acting faster than natural human response times while also reducing the effort required to stabilize in response to body pitch perturbations (Bayón et al., 2022; Beck et al., 2023). In these cases, however, the exoskeleton was ‘all knowing’ and could act instantaneously, but the real physiological system involves closed sensorimotor feedback loops. While an exoskeleton can assist or enhance motor output, it may also have undesirable impacts on the user’s biological sensory signaling (Beck et al., 2023; Canete et al., 2023). Experiments investigating sensory feedback with exoskeletons have been done but have used indirect measures of sensorimotor control such as global behavioral metrics of stability and agility or local measures of fascicle dynamics as a proxy for sensory feedback due to the difficulty of probing human proprioceptive feedback signals directly (Canete et al., 2023; King et al., 2022; Kwak and Chang, 2023). Thus, it remains unclear how exoskeletons (i.e., a stiffness in parallel with muscle-tendon units (MTU) of a lower-limb-joint) influence proprioception and performance in tasks the rely on muscle-derived sensory feedback.

Proprioceptive feedback has a large influence on motor control and stability. Muscle spindle sensory organs provide one such source of feedback, lying within the muscle belly and forming a sophisticated feedback control loop that influences motor control (Proske and Gandevia, 2012). Muscle spindle feedback has significant impact on interjoint coordination, muscle stiffness regulation, and movement and posture control and dynamics in human and animal models (Abelew et al., 2000; Gordon et al., 1995; Houk et al., 1981; Lockhart and Ting, 2007; Matthews, 1964; SAINBURG et al., 1995; Ugawa et al., 1991). Furthermore, certain indicators used to study stability and balance in humans, including within-step modulation of muscle activity, swing leg step placement during perturbations, and accurate postural and joint position control, depend on spindle feedback (Klint et al., 2008; Mayer and Akay, 2021; Sanes et al., 1985). While there is a heavy influence of sensory feedback from other avenues that are often studied in clinical applications, including visual and vestibular feedback, this previous work indicates a critical role that muscle sensory organ feedback plays in how the neuromuscular system controls movement that is independent of other sensory feedback loops. Most work quantifying this proprioceptive feedback has been limited to animal models, due to the ability to safely and effectively administer chemical agents, invasively measure afferent firing rates, isolate motor units, and highly control muscle tendon unit dynamics (Abbott et al., 2024; Abelew et al., 2000; Blum et al., 2017). These investigations have largely employed passive muscle stretches and limited active stretch paradigms without examining the influence of parallel stiffness as would be imparted by an exoskeleton (Blum et al., 2017; Honeycutt et al., 2012; Matthews, 1964; Prochazka and Gorassini, 1998; Proske and Stuart, 1985).

In part due to the limitations above, a robust model that can accurately predict the relationship between muscle spindle afferent feedback and muscle fascicle contractile dynamics in various contexts has also not yet been developed. Classically, muscle spindle feedback is believed to be influenced by muscle tendon unit (MTU) level kinematics (Kandel et al., 1991). Due to the connective tissues in parallel and series to the intra/extrafusal muscle fibers, however, whole MTU kinematics likely do not well-represent the force or length changes experienced by spindles (Blum et al., 2017; Day et al., 2017). Furthermore, afferent firing behavior is heavily dependent on the mechanical context in which the muscle is stretched and loaded, and whether the muscle is active (Blum et al., 2017; Day et al., 2017; Honeycutt et al., 2012; Matthews, 1964; Prochazka and Gorassini, 1998; Proske and Stuart, 1985). More recently, contractile force and yank (a kinetic model) acting on the intrafusal fibers has been shown to be a more accurate predictor of spindle instantaneous firing rate (IFR) than length and velocity (a kinematic model) during passive MTU stretches (Blum et al., 2017; Lin and Crago, 2002). It remains to be seen how kinetic vs. kinematic models of spindle firing perform under more complicated contractile contexts that trade off stiffness within and adjacent to an activated MTU. Regardless of the mechanism underlying spindle feedback, the altered contractile conditions resulting from exoskeleton assistance has an effect.

While exoskeletons could assist or enhance overall motor output, their effects on the internal muscle fascicle dynamics may cause unintended impacts on the proprioceptive feedback loops that are also important for accomplishing dynamic tasks. During both cyclic and balance recovery tasks, exoskeletons act mechanically in parallel to a joint, either passively through a spring in parallel with the musculoskeletal tissues, or actively through a motor coupled to the joint, they introduce an external influence on the dynamics of the MTU (Nuckols et al., 2020). By providing assistive torque, exoskeletons reduce the biological force required by muscles and increase fascicle length excursions, decreasing the biological muscle stiffness (Farris et al., 2013; Nuckols et al., 2020; Robertson and Sawicki, 2015). These effects in turn influence muscle activation dynamics and the point along the muscle’s force-length and force-velocity curves at which it operates (Robertson et al., 2017) and could offer a unique opportunity to investigate mechanisms driving spindle sensory feedback. Exoskeletons effectively disrupt the typical relationship between muscle fascicle kinematics and kinetics during active stretches. Turning up the dial on exoskeleton assistance imparts a unique context to the muscle fascicles and embedded spindle organs because as increasing exoskeleton stiffness is traded for decreased muscle stiffness to maintain overall muscle-tendon unit stiffness, fascicle force decreases while fascicle length increases. This trend is in direct opposition to the scenario during passive MTU stretch where muscle and MTU stiffness are invariant. Not only do these types of contractions allow a clearer investigation of the independent effects of muscle fascicle kinematics and kinetics on primary afferent firing, but they are also highly relevant and understudied behavioral tasks, especially as movement with human-machine systems becomes more common.

Here, we leveraged an anesthetized animal model to more directly investigate how engineered stiffness placed in parallel with a biological muscle-tendon unit (i.e., akin to a passive elastic exoskeleton on a human joint) affects proprioceptive feedback. More specifically, we imparted length and activation-controlled stretches to rat medial gastrocnemius muscle-tendon units that simulated the observed human muscle dynamics during balance recovery responses with and without exoskeleton assistance (Beck et al., 2023). Using this preparation, we examined how muscle spindle firing, reported as Instantaneous Firing Rate (IFR), within the compliant muscle-tendon unit (MTU) changed across a range of muscle stiffness-exoskeleton stiffness combinations. Based on classical kinematic models of spindle firing, we hypothesized that trading off increased exoskeleton stiffness for decreased muscle stiffness (i.e., lower muscle force and higher muscle fascicle stretch) would increase spindle IFR.

<u>Note:</u> See Supplementary Materials for simple MTU model-based predictions using kinematic vs. kinetic inputs to determine how spindle firing may change with parallel exoskeleton stiffness (Supplementary Fig. 1).

## Methods

We designed an experimental paradigm where muscle-tendon unit (MTU) length was clamped and the total force of the MTU and a selection of attached parallel springs (i.e., exoskeletons) was controlled via muscle stimulation to emulate the joint-level dynamics observed in humans during standing balance responses with an ankle exoskeleton (Beck et al., 2023). A servo motor mounted in series with the MTU applied a stereotypical ramp-stretch, and the force was matched by trading off between the muscle force (controlled via ventral root (VR) stimulation) and the exoskeleton parallel spring stiffness. We recorded changes in muscle fascicle length alongside changes in MT force with increased exoskeleton spring stiffness, while also recording the primary afferent firing from spindle organs (Fig. 1A).

**Figure 1.**
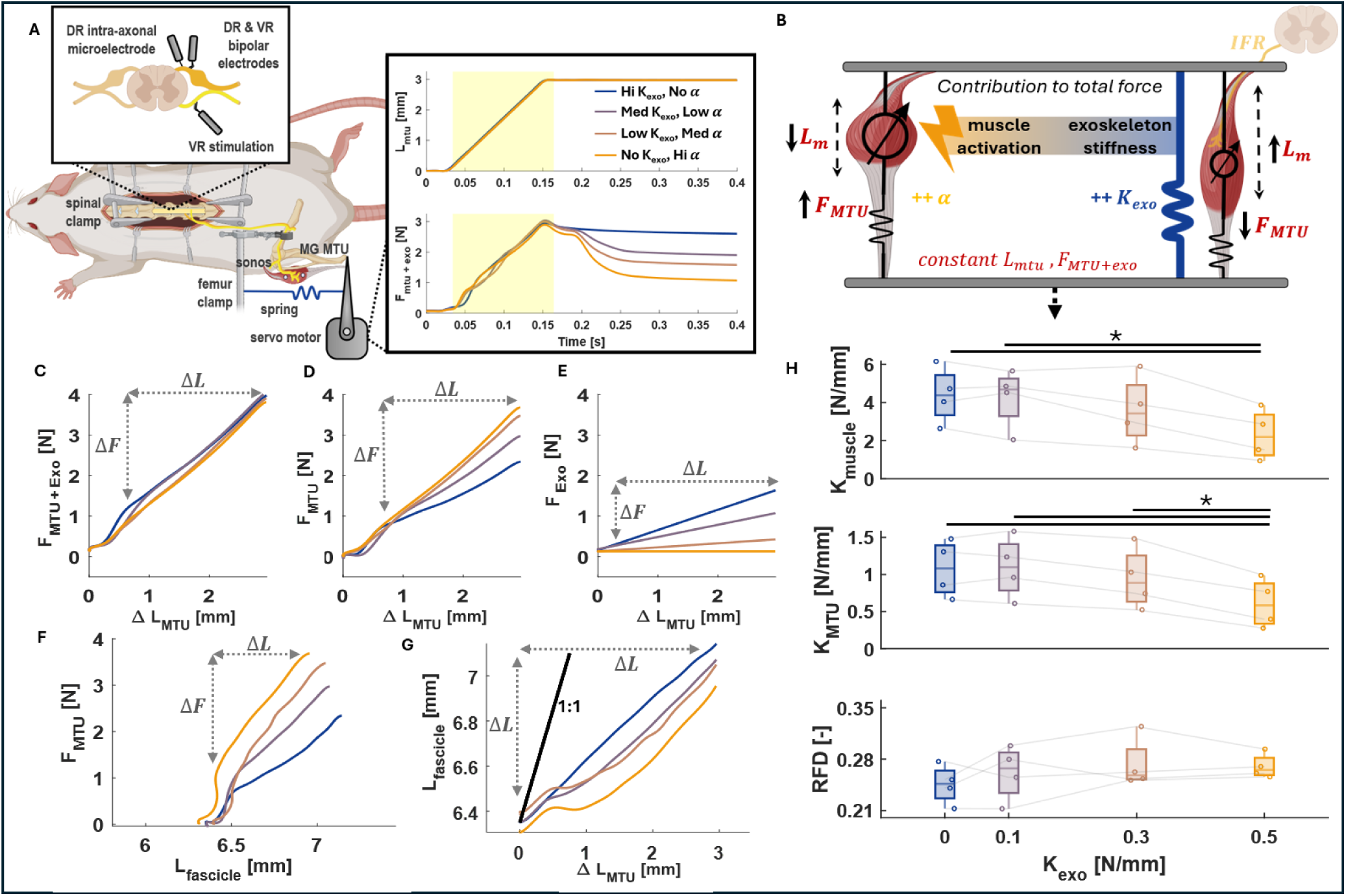
Experimental protocol. (A) Animated representation of the experimental set up. A laminectomy is performed to expose the spinal cord, and bipolar electrodes are placed on isolated dorsal (DR) and ventral root (VR) nerve fibers of interest. An intra-axonal microelectrode is inserted into the dorsal root (DR) to read afferent spikes. The femur is clamped, and the medial gastrocnemius (MG) muscle-tendon unit (MTU) is attached to a servo motor. Exoskeleton springs are fixed in parallel with the MG MTU. Muscle fascicle length changes are measured using sonomicrometry crystals (sonos). The muscle is stimulated through the ventral root (VR). Invariant total MT +exoskeleton spring force (F_mtu+exo_) is maintained during MTU stretch (L_mtu_) across all trials by using controlled electrical stimulation of the muscle during the time period indicated by the yellow shading. (B) A figurative representation showing how muscle length (L_m_) within the MT increases when reduced biological muscle-tendon (F_mtu_) force/activation (α) is traded-off for increased exoskeleton force (F_exo_)/stiffness (K_exo_) across conditions. (C-G) MTU and muscle fascicle dynamics from a representative rat. The experimentally controlled rising edge of ramp stretch derived work loops (denoted by dashed line boundaries) were investigated. (C) Total MTU + Exo force (F_mtu+exo_) and length change (ΔL_mtu_) was held invariant across exoskeleton conditions. (D-E) Relative contributions of the MTU versus Exo to total system stiffness: (D) MTU stiffness (K_mtu_) is ΔF_mtu_ divided by ΔL_mtu_ and (E) Exoskeleton stiffness (K_exo_) is ΔF_exo_ divided by ΔL_mtu_. (F) Muscle stiffness (K_muscle_) is ΔF_mtu_ divided by ΔL_fascicle_. (G) Decoupling of MTU and muscle fascicle length changes. Relative fascicle displacement (RFD) is ΔL_fascicle_ divided by ΔL_mtu_. (I) Summary data for all metrics from 4 rats (each shown by faded colored lines).

### Animal Care

All procedures and experiments were approved by the Georgia Institute of Technology Institutional Animal Care and Use Committee (protocol number A100142). The surgical methods performed were similar to those carried out in Abbott et al. (Abbott et al., 2024). Briefly, Adult female Wistar rats (250–300 g; Charles River Laboratories, Wilmington, MA, USA) were studied in terminal experiments in which they were deeply anaesthetized using isoflurane initially in an induction chamber (5% in 100% O2) and for the remainder of the experiment via a tracheal cannula (1.5−2.5% in 100% O2) while vital signs were continuously monitored. Anesthesia level and vital signs were maintained by adjusting isoflurane concentration, radiant and water-pad heat sources, and by scheduled subcutaneous injection of lactate ringer solution (1 ml/h subcutaneous).

### Data collection

IFR was measured via a glass microelectrode inserted intracellularly into the L5 dorsal root while it was suspended on bipolar hook electrodes to record the activity of single cells. Afferent firing was recorded while imparting a predetermined ramp length changes with a 3 mm amplitude, 0.125 s rise and fall time (24 mm/s rising speed), and 1 s hold using a servo motor (Aurora Scientific (Ontario, Canada)) attached to the Achilles tendon. Primary afferents were identified by their responsiveness to a 100 hz, low amplitude vibration imparted through the servo motor. The femur was clamped in place, and the Medial Gastrocnemius (MG) was isolated from the other plantar flexors, and activation of the isolated muscle is modulated with electrical stimulation imparted through the ventral root efferents. 100 Hz electrical stimulation (pulse width was 0.5ms, duration was 130ms) was imparted at amplitudes dependent on the total MTU and exoskeleton force required during the up ramp of the ramp and hold stretches. Muscle fascicle length was measured using sonomicrometry crystals embedded in the MG muscle belly (Sonometrics (Ontario, Canada)), taking care to avoid the aponeurosis. Various springs were attached in order of highest to lowest spring stiffness (ranging from 0.1 to 0.5 N/mm) in parallel to the medial gastrocnemius MTU (at the to the servo motor distally and the femur clamp proximally) to simulate a passive exoskeleton (Fig. 1A). The resulting length clamped, force-matched contractions across various exoskeleton stiffness conditions, as depicted in Fig. 1A, resembled standing balance paradigms with ramp platform perturbations with and without an exoskeleton (Beck et al., 2023). All data were collected with a Cambridge Electronic Design Power 1401 and using Spike2 software and analyzed in MATLAB.

The springs used to represent a passive exoskeleton were attached via a custom tensioning mechanism by which the preset tension in the springs could be modified to have a consistent initial exoskeleton force acting across all spring conditions. The extension springs (Mcmaster-Carr) had hook ends and were all the same initial length, so that once the tensioning mechanism was adjusted for the specific rat model’s MTU length, all springs could be easily swapped throughout the experiment to test across all experimental conditions.

### Data Processing and Analysis

Measurements of motor position, motor force and fascicle displacement lowpass filtered with a fourth-order Butterworth filter with a 50 Hz cutoff. They were then differentiated with respect to time to get the yank 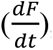, the velocity 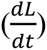, and the acceleration 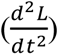. Muscle fascicle power, estimated as the muscle-tendon force multiplied by the fascicle velocity, was also analyzed as it potentially relates more directly to the dynamics experienced by the intrafusal fibers. The total MTU and exoskeleton system displacement and force were measured by the servomotor’s displacement and measured force. The exoskeleton force was calculated as the passive spring stiffness multiplied by the displacement of the spring, which was equivalent to the displacement of the total MTU and exoskeleton system. The tensioning mechanism was adjusted for each animal to ensure that the springs always start at resting length (no load) and the initial exoskeleton force is therefore zero. The biological MTU force was then calculated as the total force minus the exoskeleton force. This was further validated experimentally by comparing passive stretches of the total MTU and exoskeleton system across all spring conditions (no spring, 0.1 N-mm, 0.3 N-mm, 0.5 N-mm). This was done by subtracting the passive, no spring stretch condition from the other spring conditions and then fitting a linear function to the resulting force length curve (i.e. the stiffness of the spring in the exoskeleton’s force contribution).

To validate the effects of the parallel exoskeleton, the length change of the fascicle relative to the MTU, the stiffness of the MTU, and the stiffness of the fascicle were computed for each exoskeleton condition type. Stiffness (N/mm) was computed by fitting a linear function to force/length curves during the up-ramp of the ramp-and-hold stretches. Stiffness (in N/mm) was calculated by computing the slope of the tangent line at these force thresholds. The relative fascicle displacement (RFD) was determined in the same manner, estimating the slope 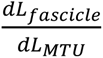 (Fig. 1G).

Spike count during the up-ramp, dynamic responses, and static responses were extracted from firing rates of the ramp-and-hold stretches (Fig. 2A-B). The dynamic response was taken as instantaneous firing rate at the peak MTU force production, when it first reaches its max stretch. The static response was taken as the firing rate approximately 0.5 s into the hold phase, and the dynamic index is the difference between the dynamic and static response.

**Figure 2:**
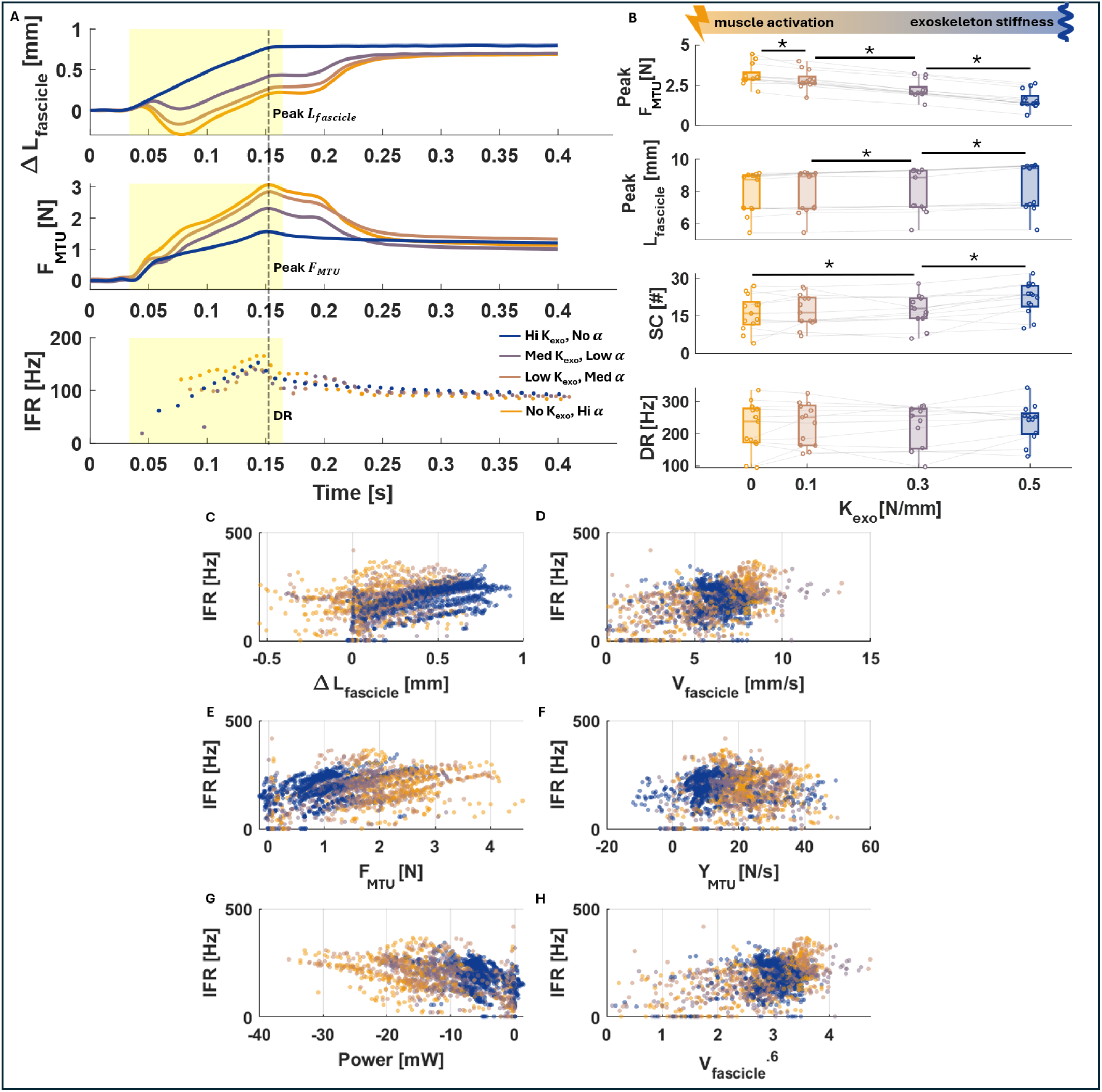
Experimental data. (A) Time series of muscle fascicle length change (ΔL_fascicle_) (top), muscle-tendon force (ΔF_mtu_) (middle) and spindle firing rate (IFR) (bottom) from a representative cell. Peak and average muscle fascicle length and MTU force varied greatly across the range of increasing exoskeleton stiffness (K_exo_) and decreasing muscle activation (α) conditions (i.e., from orange to blue), despite maintaining invariant total biological muscle-tendon plus exoskeleton force (F_mtu+exo_) across all trials during the period indicated by the yellow shading, where muscle force was controlled using electrical stimulation (α) (also see Fig. 1 A,C). The dots in the IFR traces occur at the time stamps in which a spike was recorded. (B) Summary data across all 13 cells (each represented by faded colored lines) taken from the timepoint denoted by dashed vertical line in panel A denoting peak observed muscle-tendon force (Peak F_mtu_) and muscle-length (Peak ΔL_fascicle_) during he period of active ramp-stretch. Significant changes are represented by a star and line between values and detailed model statistic are summarized in Table 1. (C-H) Correlation fits between spindle firing (IFR) and various muscle fascicle kinematic and kinetic measures across all 13 cells. Correlations were fit to the controlled portions of the ramps and ignored the initial burst during the trials in which they occurred.

**Table 1:** ANOVA model statistics with exoskeleton spring stiffness condition (K_exo_) as fixed effect.

| Variable | Degrees of Freedom | F stat | p value |
| --- | --- | --- | --- |
| $K_m$ | 3 | 21.115 | *2.1E-04 |
| $K_{mtu}$ | 3 | 43.113 | *1.2E-05 |
| RFD | 3 | 1.383 | 0.3 |
| $F_{mtu}$ | 3 | 665.003 | *1.5E-31 |
| $L_{fasc}$ | 3 | 36.662 | *4.9E-11 |
| DR | 3 | 1.005 | 0.4 |
| SC | 3 | 11.365 | *2.2E-05 |

Data were excluded from analysis if the sensory afferent action potentials could not be discriminated from the cell recording. Data were also excluded if the muscle fascicle dynamics during the trials did not reflect those expected from standing balance trials with an exoskeleton (i.e., decreased muscle force production and increased muscle stretch with increased parallel exoskeleton stiffness. See example in (Supplementary Fig. 2)). If we could not retain stable measurement from the cell long enough to record all exoskeleton conditions, it was also excluded from analysis.

Data were collected from 24 unique primary afferent cells across 8 animals and reduced to 13 cells across 4 animals after filtering based on the exclusion criteria mentioned above. In total, 117 ramp-and-hold stretches across the 4 exoskeleton conditions were analyzed. Statistical comparisons of firing metrics and muscle dynamics with exoskeleton stiffness (Fig. 2B) were conducted using repeated measures ANOVA in MATLAB. (random effect: afferent cell; fixed effect: exoskeleton spring stiffness (see Table 1)), followed by a Bonferroni post-hoc test if significance was found to determine differences within condition. Statistical comparisons of muscle stiffness, MTU stiffness, and relative fascicle displacement (RFD) with exoskeleton stiffness (Fig. 1C-G) were also conducted using repeated measures ANOVA in MATLAB. (random effect: animal; fixed effect: exoskeleton spring stiffness, (see Table 1)), followed by a Tukey’s HSD post-hoc test if significance was found to determine within condition differences. Significance was set to p<0.05. We used a Jarque–Bera two-sided goodness-of-fit test to confirm normal distribution for applicability of all ANOVAs. To study the correlation between various muscle dynamic variables (muscle fascicle length, velocity, force, yank, power, and velocity to the 0.6 power) on IFR (time series), we fitted a linear mixed-effects (LME) model with individual cells as random effects nested within each animal using a restricted maximum likelihood (REML) fit method. The models included the dynamic variable (time series) in question and the exoskeleton spring condition (categorical) as fixed effects (Fig. 2 C-H, Table 2). The LME analyses were completed in MATLAB using the fitlme function. For these analyses, initial bursts in the firing due to short range stiffness were ignored, and only the rising edge of the ramp stretch was considered. This is because that is the period during which the muscle-tendon force was controlled using muscle stimulation to match the total force and length output of the MTU + Exo system over the ramp stretch (Fig 1A, Fig. 2A – yellow shaded region). Full statistical results are reported in Tables 1 and 2. Reported values are the mean ± SE unless stated otherwise.

**Table 2:** Statistics for different LME models of spindle firing shown in Figure 2 C-H.

| Fixed Effect (95% CI) | Estimate | Standard Error | t stat | Degrees of Freedom | p value |
| --- | --- | --- | --- | --- | --- |
| $L_{fasc}$ | 199.991 | 19.865 | 10.068 | 2004 | 2.72E-23 |
| $F_{mtu}$ | 67.261 | 8.550 | 7.867 | 2004 | 5.88E-15 |
| $V_{fasc}$ | 14.217 | 1.914 | 7.430 | 2004 | 1.60E-13 |
| $Y_{mtu}$ | 0.860 | 0.301 | 2.856 | 2004 | 4.33E-03 |
| Power | -8.565 | 0.721 | -11.884 | 2004 | 1.60E-31 |
| $V_{fasc}^{0.6}$ | 44.124 | 5.625 | 7.844 | 2004 | 7.02E-15 |

## Results

In this experiment, we successfully recreated the muscle fascicle dynamics observed in human standing balance experiments with exoskeleton assistance, where the biological moment (∼muscle-tendon force) and exoskeleton torque (∼exoskeleton force) trade-off across conditions with increasing exoskeleton stiffness and decreasing muscle activation (Fig. 1A,B). The intended trade-off between muscle stiffness (K_m_) and exoskeleton stiffness (K_exo_) was achieved, indicating that the experimental set up properly recreated the *in vivo* muscle dynamics observed in humans when wearing a passive exoskeleton (Fig. 1C-G) (Nuckols et al., 2020). The measured muscle stiffness and total MTU stiffness decreased respectively from 4.38 ± 1.46 N/mm and 1.08 ± 0.38 N/mm to 2.29 ± 1.32 N/mm and 0.61 ± 0.33 N/mm as exoskeleton stiffness increased from 0 N/mm to 0.5 N/mm (p < .01, Table 1) (Fig. 1H). Overall, adding the parallel exoskeleton spring did not significantly alter fascicle displacement relative to MTU displacement during stretch (Fig. 1G). The RFD did not change significantly across exoskeleton conditions (p > .05, Table 1).

Figure 2A shows the fascicle length change (ΔL_fascicle_), biological muscle-tendon force (F_mtu_), and spindle IFR from a representative animal and cell across all exoskeleton stiffness -muscle activation conditions, with the yellow shading indicating time period of muscle stimulation. We observed a significant decrease in muscle-tendon unit (MTU) force (F_mtu_) and an increase in muscle fascicle length (ΔL_fascicle_) as exoskeleton stiffness increased. The peak absolute fascicle length increased from 7.89 ± 1.29 mm to 8.33 ± 1.45 mm (p < .0002) and the peak MTU force decreased from 3.13 ± 0.64 N to 1.59 ± 0.56 N as parallel stiffness increased from 0 N/mm to 0.5 N/mm (p < .0001,, Table 1) (Fig. 2B).

While the *in-situ* measured IFR dynamic response (DR) showed no significant change across the range of exoskeleton stiffness (p > .05), the spike count (SC) (number of spikes on the rising phase of the ramp stretch) significantly increased from 17.12 ± 7.02 to 23.03 ± 6.41 as parallel stiffness increased from 0 N/mm to 0.5 N/mm (p < .015, Table 1) (Fig. 2B). This is likely because with increased muscle activation, the fascicles shortened more initially before stretching. During this initial shortening phase, the spindle intrafusal fibers are slackened and then engage in a more rapid lengthening during the subsequent stretch, resulting in fewer overall spikes during the ramp, but reaching the same dynamic response as the relaxed condition which exhibits no initial shortening.

Spindle IFR significantly correlated with *all* muscle fascicle dynamics factors investigated, including fascicle length (β = 120.556 ± 19.865, p< 0.001, Table 2), fascicle velocity (β = 14.217 ± 1.914, p< 0.001, Table 2), muscle-tendon force (β = 67.261 ± 8.55, p< 0.001, Table 2), and muscle-tendon yank (β = 0.860 ± 0.301, p< 0.005, Table 2), during the controlled portion of the ramp stretch (Fig. 2C-F). IFR also correlated to other proposed IFR model variables (Prochazka and Gorassini, 1998), including muscle fascicle mechanical power (β = -8.565 ± 0.721, p< 0.001, Table 2) and muscle fascicle velocity raised to the power of 0.6 (β = 44.124 ± 5.625, p< 0.001, Table 2) (Fig. 2G-H). Random effects caused by differences between individual cells were significant.

## Discussion

To our knowledge, this is the first benchtop *in-situ* investigation attempting to emulate the effects of exoskeleton assistance on spindle feedback in an animal model during active contractions. Our experimental setup effectively reproduced *in vivo* muscle fascicle dynamics observed from ultrasound and electromyography studies in humans using ankle exoskeletons (Fig. 1C-G, Fig. 2A) (Beck et al., 2023; Nuckols et al., 2020). The unique mechanical context offered by parallel forces from an exoskeleton effectively decoupled muscle fascicle kinetic and kinematic factors within the MTU and enabled the opportunity to distinguish which biomechanical factors drive spindle primary afferent firing (Fig. 2C-H). As expected, trading-off increasing exoskeleton stiffness with decreasing muscle activation maintained a constant total exoskeleton + MTU force output and decreases in muscle-tendon unit (MTU) force were accompanied by increases in muscle fascicle length (Fig. 2A-B).

Interestingly, we observed no significant change in the dynamic response (DR) of spindle afferent cell IFR with exoskeleton assistance and there was no clear alignment between the DR and either the peak muscle tendon unit kinetics or muscle fascicle kinematics over the period of muscle-tendon unit stretch (Fig. 2B). Indeed, the complex interaction of increasing fascicle stretch accompanied by decreasing MTU force that is imparted by maintaining total stiffness of the exoskeleton+MTU system appeared to have a ‘cancellation’ effect on the dynamics of the underlying intrafusal muscle fibers that the primary afferent endings are wrapped around. We note that the IFR during the period of controlled ramp-stretch was significantly correlated with all kinetic and kinematic MTU and muscle fascicle dynamic variables we compared against when taken individually (Figure 2C-H, Table 2). It could be that although the intrafusal fibers containing the spindles are embedded within the extrafusal muscle fibers, the dynamics they experience are influenced by, but not directly or exclusively associated with the 1-dimensional muscle fascicle level dynamics we estimated using our force ergometer and sonomicrometry crystals. Muscle spindle deformation (and ultimately firing) may correlate with a more complex or nonlinear set of biomechanical factors than the MTU and muscle level kinematic and kinetic variables presented here (Blum et al., 2020, 2017; Matthews and Stein, 1969).

There have been numerous attempts to accurately represent spindle firing and the dynamics of the intrafusal muscle fibers which the primary afferent endings are wrapped around (Proske and Gandevia, 2012). Classical approaches involve investigating different MTU and fascicle states and their correlation with muscle spindle firing (Prochazka and Gorassini, 1998), but our results suggest that none of these proposed models can fully account for the firing dynamics we measured in response to the combination of reduced MTU force and increased muscle fascicle stretch imparted by parallel exoskeleton stiffness (Fig. 3, Table 3). Recently, Blum et al. showed that kinetic models outperform kinematic models in predicting IFR generated during passive stretches *in-situ*. IFR predictions using Blum’s models with a simple muscle and exoskeleton model are described in the Supplementary Materials (Supp. Fig. 1C). Attempts to fit such models directly to our experimental data revealed that *neither* the kinetic *nor* kinematic models could reliably predict spindle firing in our data across exoskeleton conditions (Fig. 3A-B, Table 3) (Blum et al., 2017).

**Figure 3:**
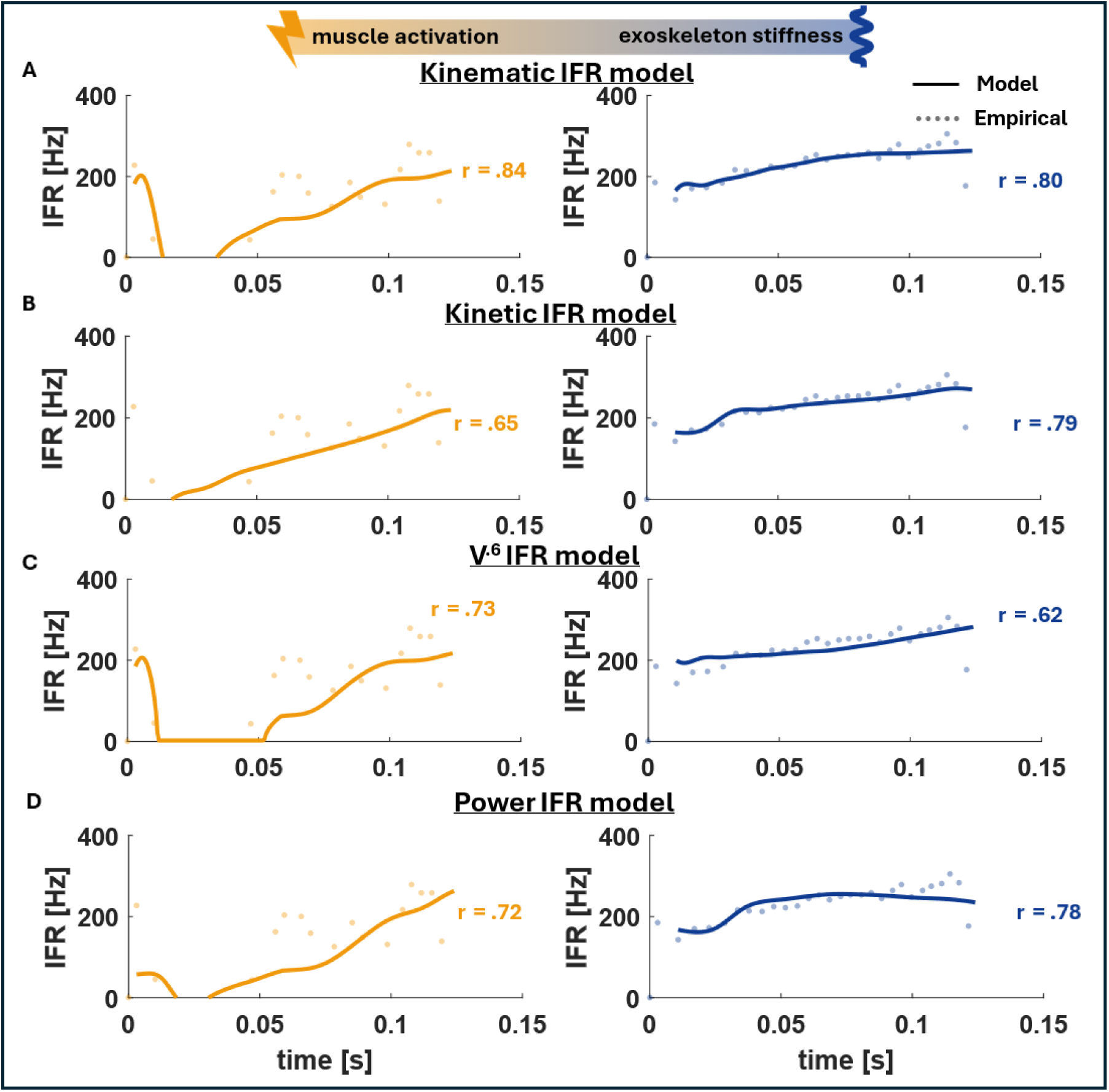
Time-series of spindle firing (IFR) from a representative cell within the same rat model, overlaid with fits from three firing models described in previous literature. (Left) Zero exoskeleton stiffness (K_exo_ **=0**), high muscle activation condition. (Right) High exoskeleton stiffness (K_exo_ **=** max), no muscle activation (=relaxed) condition. A) The Kinematic IFR model is represented by the equation *IFR_L_* = (Δ*L_m_* + *b_L_*)*k_L_* + (*V_m_* + *b_V_*)*k_V_* (Blum et al., 2017). B) The Kinetic IFR model is represented by the equation *IFR_F_* = (*F_c_* + *b_F_*)*k_F_* + (*Y_c_* + *b_Y_*)*k_Y_* (Blum et al., 2017). C) The V^0.6^ model is represented by the equation *IFR_v.6_* = (*V*^0.6^ + *b_v_*)*k_v_* (Prochazka and Gorassini, 1998). D) The Power IFR model is represented by the equation *IFR_power_* = (*Power* + *b_p_*)*k_p_*. Full details of fit equations can be found in Table 3.

**Table 3:** IFR model fit parameters from data in Figure 3.

| IFR Model | $K_{\text{exo}} = 0$ condition fit equation | $K_{\text{exo}} = 0$<br>$R^2$ | $K_{\text{exo}} = \text{max}$ condition fit equation | $K_{\text{exo}} = \text{max}$<br>$R^2$ |
| --- | --- | --- | --- | --- |
| Kinetic Model | $43.47 * (F_{\text{mtu}} + 2.14) - (3.74 * 10^{-12}) * (Y_{\text{mtu}} - 61.05)$ | 0.57 | $33.21 * (F_{\text{mtu}} + 5.35) - 4.61 * (Y_{\text{mtu}} - 17.93)$ | 0.85 |
| Kinematic Model | $(2.16 \times 10^{-10}) * (L_{fasc} + 20.89) - 14.68 * (V_{fasc} + 9.01)$ | 0.57 | $104.1 * (L_{fasc} + .12) - 25.4 * (V_{fasc} .66)$ | 0.84 |
| $V_{fasc}^{0.6}$ Model | $31.56 * (V_{fasc}^{0.6} + 3.94)$ | 0.45 | $38.33 * (V_{fasc}^{0.6} + 2.94)$ | 0.57 |
| Power Model | $-5.94 * (Power + 22.49)$ | 0.47 | $-24.35 * (Power - 4.82)$ | 0.85 |

One approach that may shed some light on the results in this experiment is to consider non-linear or viscoelastic models of the intrafusal and extrafusal muscle fibers to predict primary and secondary afferent firing (Corominas-Murtra and Petridou, 2021; Mileusnic et al., 2006; Rehorn et al., 2014). A nonlinear velocity model (V^0.6^) performed similarly to other models we examined, although it did not capture some of the IFR dynamics that existed across exoskeleton condition (Fig. 3C). We also examined muscle fascicle mechanical power, which depends on a combination of kinetic and kinematic factors - both extrafusal fascicle force and velocity. We surmised that mechanical power which captures the rate of change in material strain energy might be a quantity that could translate more directly to bulk local 3D dynamics experienced by the intrafusal fibers. In addition, other studies have pointed toward the potential that spindles encode more abstract energy/effort based representations of biomechanical state during locomotion. For example, Kwak et al. suggested that the fascicle dynamics sensed by the muscle spindles could reflect the whole-body walking economy (Kwak and Chang, 2023). Furthermore, Monjo and Allen suggested more recently that afferent signals from muscle spindles may play a significant role in effort perception, suggesting that the brain receives information about the strength of fusimotor commands through an increase in afferent firing rate driven by gamma drives (Monjo and Allen, 2023). Despite our rationale for mechanical power as a potential driving factor of spindle firing, similar to other models, power also yielded similarly mixed results in predicting spindle firing across exoskeleton conditions (Fig. 3D).

Our finding that spindle firing does not change when applying the muscle-level neuromechanical conditions observed *in vivo* during walking and balancing with lower-limb exoskeletons lends some support for the idea that sensorimotor adaptation in this context must occur from other mechanisms driven by descending commands, such as modulating gamma drive. Previous work has demonstrated significant sensorimotor adaptation to exoskeleton assistance (Kao et al., 2010a, 2010b; Steele et al., 2017), but few studies have attempted to identify the underlying neural mechanisms. For example, Kao et al. (Kao et al., 2010a), demonstrated a 36% reduction in soleus EMG in humans walking with a powered ankle exoskeleton and in a follow up study (Kao et al., 2010b) showed that soleus H-reflex modulation remains unaltered with powered exoskeleton assistance in unimpaired human users (Kao et al., 2010b, 2010a). This indicates that increased presynaptic inhibition of Ia afferents is likely not the mechanism by which soleus EMG is reduced with exoskeleton assistance. Although we found that changes in neuromechanical context due to exoskeleton assistive force had little effect on primary afferent feedback, a limitation of our study is that gamma drive is inhibited in these preparations. Therefore, the afferent feedback measured from the spindle fibers will not be the same as when the spindles are tuned via gamma activation. It is possible that the altered body-environment dynamics introduced by exoskeletons impact gamma drive in human users, and this could act to modify the output from spindle feedback. In addition, changes in motor output resulting from donning exoskeletons may result from modulation in higher-level cortical control that elicits a more direct attenuation of supra-spinal drive, especially in the early stages of learning the novel task (Steele et al., 2017).

Intrafusal muscle fibers have contractile dynamics independent of the extrafusal fibers they are embedded within. Muscle spindle firing, in turn, results from mechanotransduction of the muscle spindle group primary afferents in the nerve endings wrapped around these intrafusal muscle fibers. Understanding the interaction between these fiber types may, therefore, be necessary to explain the changes in muscle spindle firing due to an added exoskeleton in parallel to the MTU influencing extrafusal muscle fiber dynamics. Our recent biophysical muscle spindle model is a potential candidate for this (Blum et al., 2020; Simha and Ting, 2024). The model simulates extrafusal muscle fiber length changes based on the experimentally measured motor forces and applies this length to the two types of intrafusal fibers, bag 1 and chain, under the assumption that intrafusal fibers act in parallel with the extrafusal fiber. All muscle forces are modelled based on cross-bridge dynamics, with each fiber type having different properties of crossbridge dynamics. Each muscle fiber also receives its own independent motor drive—alpha drive to the extrafusal muscle fiber, gamma static to the chain fiber, and gamma dynamic to the bag 1 fiber. The model also applies a normalized and scaled value of alpha activation that was used in the experiment and kept the gamma motor activation at a minimal level to simulate alpha activation below the gamma threshold. This model may, thereby, allow us to more accurately predict muscle spindle firing by simulating the dynamics that the intrafusal muscle fibers experience directly.

This study provides initial insight along a path to mechanistic understanding of how wearable robotic systems impact sensorimotor integration during unsteady dynamic movements that require sensory feedback control. Here, we successfully replicated muscle-level dynamics observed in human ankle exoskeleton studies using a benchtop in-situ rat preparation by placing an elastic exoskeleton in parallel with the exposed medial gastrocnemius muscle tendon unit. Trading-off increasing exoskeleton stiffness with decreasing muscle stiffness drastically altered muscle fascicle force and length dynamics but did not significantly alter muscle spindle Instantaneous Firing Rate (IFR). This finding is critical for informing future muscle spindle feedback models and suggests that more complex biophysical models of intrafusal fiber dynamics must be developed. Furthermore, these findings imply that the nervous system’s adaptation to exoskeleton assistance may rely on higher-order modulation, such as changes in gamma drive or supraspinal control, rather than direct alterations in peripheral spindle sensitivity. Next experiments could employ decerebrate animal preparations to enable the study of exoskeleton interaction with in-vivo systems with intact spinal and peripheral circuitry but without higher cortical interference. This could provide a powerful framework to isolate impact of exoskeletons on reflexive and gamma-mediated control mechanisms. Complementary experiments using microneurography in humans could help extend these insights to understand if and how exoskeletons could augment human stability or agility during sensorimotor control tasks requiring rapid response to unsteady conditions and whether humans can adapt to robotic assistance over longer timescales to become super-human movers.

## Acknowledgments

We would like to thank Jake Stephens for technical support and feedback during this project.

