## Supplementary Figures for "Emulating the influence of exoskeleton stiffness on primary afferent feedback in rat isolated muscle-tendon unit"

### Supplementary Materials

We developed a simple (neuro)mechanical model to give insight into how an elastic exoskeleton might influence spindle afferent firing during ramp stretches across a range of muscle activation - exoskeleton stiffness combinations that maintained a total constant system (=muscle-tendon unit (MTU) + exoskeleton (Exo)) stiffness (Supp. Fig. 1). Our muscle-tendon unit (MTU) model builds on that developed by Blum et al., in which a muscle and muscle spindle are represented in parallel and then placed in series with a tendon (Blum et al., 2020). Following Blum, we split the muscle into its contractile (C) and non-contractile (NC) element and added a tendon ($K_t$) in series with the muscle to make a muscle-tendon unit (MTU). Then, in addition, we added a passive exoskeleton spring ($K_{exo}$) in parallel with the whole MTU (Supp. Fig. 1A). Muscle-tendon length changes ($L_{mtu}$) were applied to the model while modulating active stiffness of the muscle ($K_m$) to maintain a constant total force output ($F_{total}$), that includes the biological muscle-tendon force ($F_{mtu}$) and the exoskeleton force ($F_{exo}$). Elastic exoskeletons of varying stiffness were added in parallel to the MTU to determine the effect on the predicted spindle IFR. Force in the noncontractile component ($F_{nc}$) was determined according to (Blum et al., 2020) $F_{nc} = K_{lin}(\Delta L_m) + Ae^{K_{exp}(\Delta L_m)}$, where $\Delta L_m$ approximates the muscle fascicle length change, and $K_{lin}$, $K_{exp}$, and A are optimized fit constants. $F_c$ was then determined by subtracting $F_{nc}$ from $F_{mtu}$ (as $F_{mtu} = F_c + F_{nc}$). Then, continuing to follow Blum, $F_c$ was used to predict IFR in a kinetic based IFR model: $IFR_f = (F_c + B_f)K_f + (Y_c + B_y)K_y + C$, where $Y_c$ is the Yank, defined as the time derivative of $F_c$, and $B_f$, $K_f$, $B_y$, $K_y$, C are optimized fit constants. Similarly, the muscle fascicle length change ($\Delta L_m$) and velocity ($V_m$) were used to predict IFR in a kinematic-based IFR model: $IFR_l = (\Delta L_m + B_l)K_l + (V_m + B_v)K_v + C$.

To understand how the trade-off between exoskeleton ($K_{exo}$) and active muscle stiffness ($K_m$) would decouple muscle fascicle length and force during ramp stretches, we chose to represent the system in terms of its component stiffnesses $K_{mtu}$, $K_m$, $K_t$ and $K_{exo}$ (see Eqs. 1-4). With an added exoskeleton stiffness ($K_{exo}$) in parallel to the MTU, the total force is split between the exoskeleton and the MTU, resulting in the following stiffness characteristics.

$$1)\quad K_{mtu+exo} = \frac{F_{mtu}+F_{exo}}{L_{mtu}}$$

$$2)\quad K_{mtu} = \frac{K_m K_t}{K_m+K_t}$$

$$3)\quad K_{mtu+exo} = \frac{K_{m\,(w/exo)}K_t}{K_{m\,(w/exo)}+K_t} + K_{exo}$$

$$4)\quad K_{m\,(w/exo)} = \frac{K_t(K_{mtu+exo} - K_{exo})}{K_t + K_{exo} + K_{mtu+exo}}$$

With $K_t$ (tendon stiffness) and $K_{mtu+exo}$ as constants, equation (4) shows that as exoskeleton stiffness ($K_{exo}$) increases, muscle stiffness ($K_m = \frac{F_{mtu}}{\Delta L_m}$) decreases, and vice versa (Supp. Fig. 1D). As a result, IFR decreases with increasing parallel exoskeleton stiffness when determined as a function of muscle fascicle kinetics; but IFR increases with increasing parallel exoskeleton stiffness when determined as a function of muscle fascicle kinematics (Supp. Fig. 1C). The *in-situ* experiments detailed in this study were designed to help resolve the contrasting predictions from classical kinematic vs. more recent kinetic-based models for spindle output IFR during ramp stretches of a MTU that trade-off active muscle stiffness $K_m$ and parallel exoskeleton stiffness $K_{exo}$.

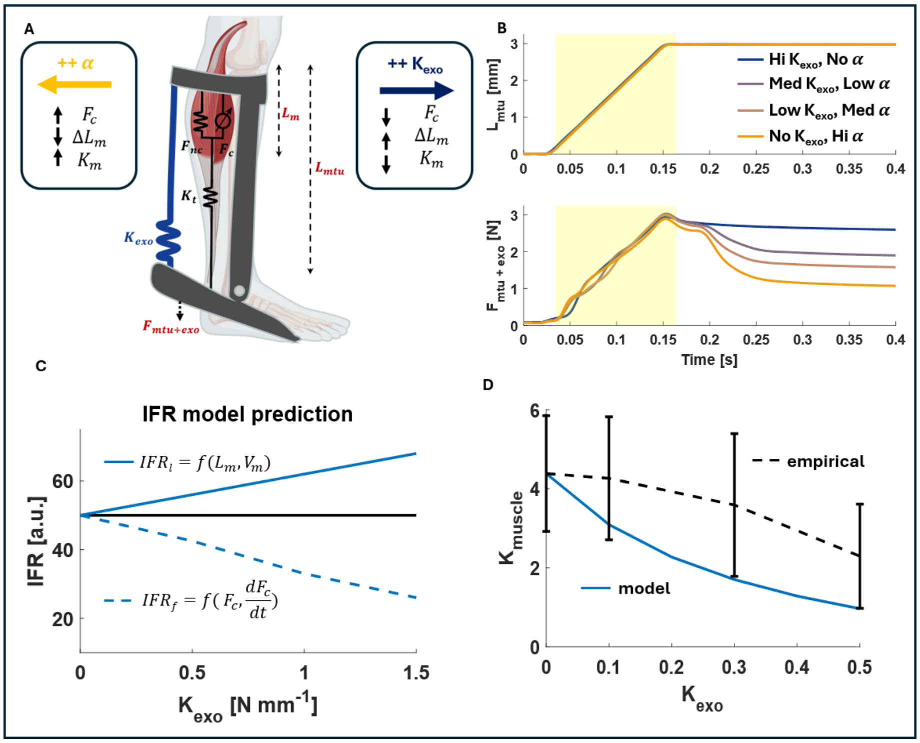


**Supplementary Figure 1:** (A) A simple Hill-type muscle-tendon model with contractile ($F_c$) and non-contractile ($F_{nc}$) muscle forces contributing to the total biological muscle-tendon force ($F_{mtu}$), plus a tendon ($K_t$) included as a series elastic element, and the addition of an 'exoskeleton' ($K_{exo}$) as an elastic element in parallel with the whole MTU. Muscle length ($L_m$) was defined as the difference between the muscle-tendon length ($L_{mtu}$) and the tendon length ($L_t$). (B) In order to predict spindle firing in the context of an elastic exoskeleton, we applied ramp-stretches to the model. To match *in vivo* observations from analogous human experiments and the *in situ* rat protocol we executed in this study, external $F_{mtu+exo}$ and $L_{mtu}$ dynamics were held invariant across all conditions by trading-off contributions of muscle stiffness, $K_{muscle}$ (via activation α) and exoskeleton stiffness, $K_{exo}$ to the total $F_{mtu+exo}$ force during MTU stretch. The time during which the muscle was 'activated' is depicted by the yellow shading. (C) Model predicted spindle afferent firing pattens based on a kinetic ($IFR_f$) (dashed blue) versus a classical kinematic ($IFR_l$) (dark blue) model of muscle spindle firing. (D) Model prediction of muscle stiffness ($K_{muscle}$) versus exoskeleton stiffness ($K_{exo}$) compared to empirical observations in the current *in situ* rat study. Empirical muscle stiffness ($K_{muscle}$) was estimated by dividing the change in muscle-tendon force ($\Delta F_{mtu}$) (assuming $F_{mtu}$ ~= to $F_m = F_c + Fnc$) by the change in muscle fascicle length ($\Delta L_m$) recorded using sonomicrometry. Error bars shown are ±SEM.

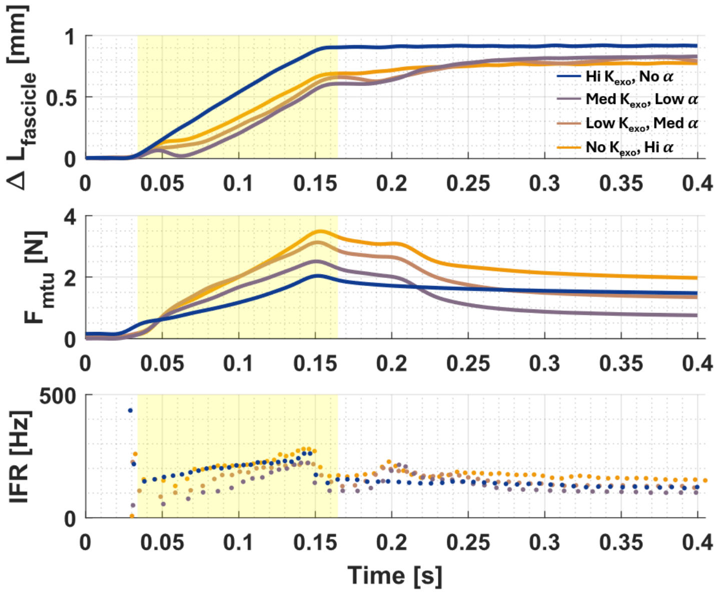


**Supplementary Figure 2:** Example of an excluded trial in which muscle fascicle dynamics did not reflect those expected based on *in vivo* ultrasound recordings during standing balance trials from humans using ankle exoskeletons (e.g., in Beck et al. (2023). In the cases observed on Beck et al., the smallest muscle fascicle stretch occurred without the exoskeleton (orange) and increasing exoskeleton assistance (purple to blue) resulted in decreased MTU force production and increased muscle fascicle stretch. As seen here, in trials we excluded, the no exoskeleton condition (orange) did not elicit the smallest fascicle length change as expected.
